# SCOIGET: Predicting Spatial Tumor Evolution Pattern by Inferring Spatial Copy Number Variation Distributions

**DOI:** 10.1101/2025.02.22.639602

**Authors:** Yujia Zhang, Yitao Yang, Yan Kong, Bingxu Zhong, Kenta Nakai, Hui Lu

## Abstract

Constructing a comprehensive spatiotemporal map of tumor heterogeneity is essential for understanding tumor evolution, with copy number variation (CNV) as a significant feature. Existing studies often rely on tools originally developed for single-cell data, which fail to utilize spatial information. Our study aims to develop a model that fully leverages spatial omics data to elucidate spatio-temporal changes in tumor evolution. Here, we introduce SCOIGET (Spatial COpy number Inference by Graph on Evolution of Tumor), a novel framework using graph neural networks with graph attention layers to learn spatial neighborhood features of gene expression and infer copy number variations. This approach integrates spatial multi-omics features to create a comprehensive spatial map of tumor heterogeneity. Notably, SCOIGET achieves a substantial reduction in error metrics (e.g., mean squared error, cosine similarity, and distance measures) and produces superior clustering performance as indicated by higher Silhouette Scores compared to existing methods. Our model significantly enhances the efficiency and accuracy of tumor evolution depiction, capturing detailed spatial and temporal changes within the tumor microenvironment. It is versatile and applicable to various downstream tasks, demonstrating strong generalizability across different spatial omics technologies and cancer types. This robust performance improves research efficiency and provides valuable insights into tumor progression. In conclusion, SCOIGET offers an innovative solution by integrating multiple features and advanced algorithms, providing a detailed and accurate representation of tumor heterogeneity and evolution, aiding in the development of personalized cancer treatment strategies.

**Key Points:** *1. Innovative Framework for Spatially Contextualized CNV Inference:* Introduces SCOIGET, a graph-based model that integrates spatial omics data with graph neural networks and attention mechanisms to accurately infer spatial copy number features.

*2. Detailed Tumor Heterogeneity Mapping:* Constructs representations of tumor evolution, capturing heterogeneity and clonal dynamics at both cellular and subcellular resolutions.

*3. Versatile and Robust Performance:* Outperforms existing methods across multiple datasets, demonstrating reliability and adaptability to diverse spatial omics platforms and cancer types.

*4. Broad Downstream Applications:* Enables tasks such as tumor subclone identification, clustering, and evolutionary trajectory analysis, facilitating precision oncology strategies.

## Introduction

Tumor evolution, also known as carcinogenesis, is a complex, multi-step process characterized by dynamic changes in morphology, genetic composition, and epigenetic profiles[1]. Studies have demonstrated significant differences in genetic homogeneity and heterogeneity within tumors, with pronounced variations observed between malignant regions and adjacent benign tissues[2,3]. Copy number variations (CNVs), which refer to the gains or losses of specific genomic regions, are crucial for evaluating such heterogeneity and play vital roles in influencing cell proliferation, therapeutic response, and resistance mechanisms[4,5]. Traditionally, whole-genome and whole-exome sequencing have been the primary approaches for identifying CNVs, providing a broad overview of genomic alterations based on next-generation sequencing (NGS) data[6–8]. However, these “bulk” approaches fail to capture intratumoral heterogeneity, particularly spatial heterogeneity, which is crucial for understanding tumor progression and developing targeted therapies. Although spatially resolved DNA sequencing could theoretically address this gap[9], its practical application is limited by the scarcity and high cost of such data.

Recent studies have revealed a strong correlation between CNVs and differential gene expression at the RNA level[10], suggesting that CNVs can be inferred from transcriptomic data as an alternative to direct DNA sequencing. Consequently, various algorithms have been proposed to infer CNVs from single-cell RNA sequencing (scRNA-seq) data. For instance, InferCNV[11] identifies chromosome-level CNVs by comparing the gene expression profiles of target cells to a reference population of normal cells, but its reliance on well-defined normal references limits its applicability. CopyKAT[12] and SCEVAN[13] address this issue by using unsupervised clustering to distinguish tumor cells from normal cells, but their probabilistic frameworks are constrained when handling large or highly heterogeneous datasets. CopyVAE[14], which applies a variational autoencoder (VAE) to capture nonlinear gene expression patterns for CNV inference, depends heavily on the accurate identification of diploid cells, which can be challenging in diverse tumor contexts. Furthermore, all these scRNA-seq-based methods overlook spatial information, making them inadequate for addressing spatial heterogeneity within tumors. This limitation prevents these approaches from effectively modeling the complex spatial dynamics that are critical for understanding tumor evolution.

Recognizing the insufficiency of spatial information in existing methods, studies in spatial transcriptomics (ST) have underscored the indispensable role of spatial context in accurately mapping gene expression patterns[15,16]. The recently proposed CalicoST[17] method utilizes hidden Markov models (HMM) to detect correlations within genomic intervals and hidden Markov random fields (HMRF) to model spatial correlations between cancer clones. While this approach integrates spatial data, it assumes clone state consistency across neighboring spatial regions, a strong constraint that can lead to inaccurate predictions in tumors with high spatial heterogeneity. Additionally, for low-coverage ST data, CalicoST aggregates neighboring spot counts to improve robustness, but this sacrifices spatial resolution and limits its generalizability across different platforms. Moreover, the reliance on computationally intensive statistical optimization algorithms, such as negative binomial and beta-binomial distributions, results in lengthy runtimes (2–8 hours), further constraining its scalability and applicability in large-scale studies.

To address these limitations, we propose SCOIGET (Spatial COpy number Inference by Graph on Evolution of Tumor), a novel framework that leverages spatial omics data to achieve accurate and efficient CNV inference. The key innovation of SCOIGET lies in its use of graph structures to dynamically model spatial relationships, enabling the method to capture local spatial heterogeneity and adapt to tumors with varying complexity. By incorporating graph neural networks (GNN) with graph attention (GAT) mechanisms, SCOIGET can flexibly adjust the influence of neighboring spots, overcoming the rigidity of uniform neighbor treatment. Additionally, its graph-based design allows information from low-coverage regions to propagate through adjacency relationships, minimizing the need for spot aggregation while preserving fine spatial resolution. These features ensure the robust performance of our model across diverse tumor samples and sequencing platforms. Unlike computationally intensive statistical optimizations, SCOIGET employs deep learning strategies, offering high computational efficiency and scalability for large datasets. This comprehensive integration of spatial and transcriptomic data, coupled with its adaptability and efficiency, makes SCOIGET a powerful tool for exploring tumor evolution, uncovering tumor clone dynamics, and supporting early therapeutic interventions.

## Materials and methods

### Dataset Information

We utilized four spatial transcriptomics datasets, including three colorectal cancer (CRC) cohorts and one prostate cancer (PCa) cohort. The first CRC Visium dataset was obtained from the HTAN-WUSTL atlas[18], accompanied by whole-exome sequencing (WES) data used to construct the gold standard for copy number analysis. The second dataset was derived from a *Cell* publication in 2023[19], consisting of four CRC Visium samples from the same patient covering different stages. The third dataset, sourced from the *10x Genomics* repository[20], included three CRC Visium HD samples[7]. Finally, the fourth dataset was a prostate cancer Visium dataset from a 2022 *Nature* study[21], comprising four samples collected from a single patient.

The first dataset served as a validation set, and the remaining three were used as case studies to explore the utility and generalizability of our method. Further details are provided in Supplementary Note 1 and Supplementary Table 1.

### Data Preprocessing

All datasets, generated using the 10x Genomics Visium[22] and Visium HD[23] platforms, were processed through a standardized pipeline. Spatial transcriptomics data, including gene expression matrices, spatial coordinates, and high-resolution tissue images, were integrated into AnnData objects using Python Scanpy library[24] and the BMAP platform[25]. Quality control involved filtering out cells with fewer than ten total counts and genes expressed in fewer than five spots. Cell cycle-related genes, mitochondrial genes, and human leukocyte antigen (HLA) genes were removed to minimize non-tumor-related variability. Gene expression counts were normalized to ensure a consistent total count per cell, log-transformed to stabilize variance, and scaled to reduce the influence of outliers. Diagnostic visualizations, such as histograms of QC metrics, UMAP embeddings, and spatial scatter plots, were generated to assess data integrity and spatial distributions.

### Sample integration

To integrate samples from different stages into a unified analysis, we employed the Harmony algorithm[26] to eliminate batch effects. Using the Python implementation of Harmony, harmonypy, we integrated ST data stored in an AnnData object. This process was conducted after PCA computation and before constructing the neighbor graph, ensuring effective removal of batch effects while preserving biological variation. The resulting batch-corrected embedding was used for downstream analyses, providing a harmonized dataset with minimal technical biases.

### Gene Annotation and Binning

To map gene expression data to specific genomic regions, we annotated each gene using the Ensembl genome database (Release 98). Each gene was linked to its chromosome and genomic coordinates (start and end positions) based on its Ensembl gene ID. Genes lacking valid chromosomal information were excluded to ensure data accuracy.

To reduce data sparsity and improve CNV detection, a gene binning strategy was employed. Genes were first sorted by chromosome and genomic position, then grouped into bins of 25 adjacent genes. For chromosomes where the total number of genes was not divisible by 25, lowly expressed genes were excluded to align the count. Within each bin, gene expression values were aggregated by summing, producing a condensed dataset that preserves critical genomic information while reducing noise and dimensionality.

### Spatial graph construction

To integrate spatial relationships with gene expression data, we constructed spatial neighbor graphs for each phase, iteratively refining both spatial and genomic features.

#### Initial Graph Construction

In the first phase, a k-nearest neighbors (k-NN)[27] graph was built using binned gene expression data and spatial coordinates. Each spot was connected to its five nearest neighbors based on spatial Euclidean distance, forming the graph’s adjacency matrix. Edge weights were computed by combining spatial proximity and expression similarity. Binned expression data were standardized and reduced via principal component analysis (PCA) to 32 dimensions to extract informative features. In this step, the k-NN connections were established based on PCA-transformed Euclidean distances in the feature space. Softmax normalization was then applied to convert these distances into edge probabilities, emphasizing stronger connections between nodes with similar expression profiles while incorporating spatial neighborhood information.

#### Refined Graph Construction

In the second phase, pseudo-CNV estimates from the initial round of training were incorporated as new node features, augmenting the graph with genomic information. These updated features, which combined spatial and refined genomic characteristics, were furthre standardized and reduced via PCA to 32 dimensions. A new k-NN graph was constructed using the PCA-transformed feautre space, ensuring that edge connections reflected both spatial proximity and genomic feature similarity. This iterative refinement progressively enhanced the spatial representation, improving CNV localization accuracy and spatial clone identification precision. The edge weights in this refined graph were recalculated using the updated feature embeddings, and softmax normalization was reapplied to ensure probabilistic interpretation while maintaining biologically relevant connectivity patterns.

### SCOIGET framework

The core of the SCOIGET framework is a graph neural network (GNN)[28] with graph attention layers, which specifically designed to capture complex spatial-transcriptomic-genomic interactions within spatial transcriptomics data. The model integrates gene expression profiles with spatial information to detect CNVs and infer tumor heterogeneity.

#### Input Data

SCOIGET utilizes three main components: (1) Node Features, where each spot or cell is represented by either binned gene expression data (feat) or pseudo-copy number profiles (norm_x) depending on the training stage; (2) Spatial Graph Structure, which encodes spatial relationships in an adjacency matrix (graph_neigh) where nodes represent spots or cells and edges indicate spatial proximity based on tissue architecture; and (3) Edge Attributes, which quantify the similarity between neighboring spots by calculating gene expression distances and normalizing them using softmax.

#### Model Architecture

SCOIGET’s model comprises three primary components: an Encoder, a Decoder, and a Copy Number Encoder (CNEncoder). The Encoder utilizes three Graph Attention Network (GAT) layers[29] with multiple attention heads and ReLU activations to learn latent representations that capture complex spatial-transcriptomic-genomic interactions. The Decoder reconstructs the original input features from these latent representations through fully connected layers, including an intermediate layer with 128 units and ReLU activations to preserve intricate patterns. The CNEncoder estimates CNVs from the reconstructed features by employing a Hidden Markov Model (HMM)[30], which models genomic bins with discrete states and Gaussian emission probabilities. This encoder integrates spatial smoothing to enhance CNV localization and includes a regularization loss term to prevent overfitting.

#### Copy Number Estimation and Refinement

The CNEncoder in SCOIGET estimates CNVs by identifying regions with consistent copy number states through a Hidden Markov Model (HMM). This process involves predicting hidden states that correspond to different copy number levels and applying spatial smoothing to reduce noise and improve CNV localization. The final CNV estimates are normalized and rescaled to ensure consistency across samples. Detailed implementation and parameter settings of the HMM are presented in Supplementary Note 3.

#### Training Procedure

The model undergoes two phases of training. In the first phase, it uses binned gene expression data and a spatial graph constructed from the original features. The model is trained without a validation set to learn initial latent representations and reconstruct the input features. Following this phase, the CNEncoder estimates pseudo-copy numbers using a HMM, generating initial CNV predictions. In the second phase, the pseudo-copy numbers are incorporated into the node features, and a new spatial graph is created. The model is retrained on this updated graph, utilizing a validation set to monitor performance and prevent overfitting, with early stopping based on validation loss. The CNEncoder re-estimates the copy numbers, refining the CNV profiles with the updated model.

Details of the model can be found in Supplementary Note 2.

### Loss Function

The overall loss function includes several components to ensure effective training.

***Reconstruction Loss:*** measures the discrepancy between the original and reconstructed features, encouraging the model to retain essential information in the latent space:

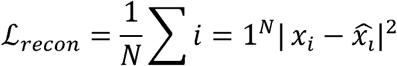

***KL Divergence Loss:*** regularizes the latent space by encouraging the learned distribution to be close to a prior distribution, which prevents overfitting and ensures meaningful latent representations:

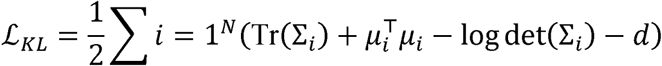

where *μ_i_* and Σ*_i_* are the mean and covariance of the latent distribution for sample *i*, and *d* is the dimensionality of the latent space.

***Regularization Loss:*** uses L2 regularization on the reconstructed features helps prevent overfitting and encourages smooth predictions:

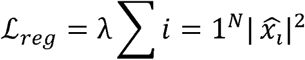

where *λ* controls the contribution of the regularization term.

***Spatial Smoothing Loss:*** enforces spatial consistency in the predicted copy numbers by minimizing the discrepancy between the copy numbers of connected nodes in the graph:

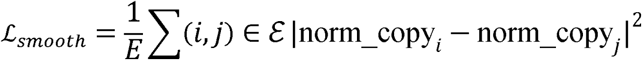

where *ε* is the set of edges in the graph, and *E* is the number of edges.

The ***Total Loss*** is a weighted sum of the reconstruction loss, KL divergence loss, regularization loss, and spatial smoothing terms:

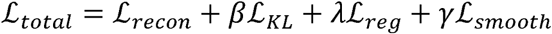

### Model Implementation and Training

The model is implemented using PyTorch and PyTorch Geometric (PyG)[31] libraries, leveraging GPU acceleration for improved performance. Training is conducted with the Adam optimizer, typically set with a learning rate of 0.001. During each epoch, the model performs forward propagation to compute outputs and loss, followed by backpropagation to update the model parameters. In the first training phase, the focus is on learning meaningful latent representations without the use of a validation set. In the second phase, a validation set (20% of the data) is introduced to monitor model performance and guide training adjustments, with early stopping employed if the validation loss does not improve.

### Baseline Methods

To evaluate the performance of SCOIGET, we compare it with four existing CNV inference algorithms: InferCNV[11], CopyVAE[14], CopyKAT[12], and SCEVAN[13]. InferCNV estimates CNVs by comparing gene expression levels between tumor and reference normal cells, employing a sliding window approach to smooth the expression signals. CopyVAE, a variational autoencoder-based method, infers CNVs from single-cell RNA-seq data, capturing the probabilistic nature of CNV states. CopyKAT segments tumor cells based on gene expression levels to identify large-scale CNVs, effectively distinguishing tumor cells from normal ones. SCEVAN integrates spatial and genomic information to infer CNVs, leveraging advanced statistical models to improve detection accuracy. Since CalicoST requires raw sequencing files for computation, we did not include it in the comparison here due to data availability constraints. All algorithms are executed using their recommended default parameters, as specified in the original publications or software repositories. WES data is used as the baseline for copy number estimation accuracy.

### Evaluation Metrics

The performance of SCOIGET and the contrast algorithms is comprehensively evaluated using several metrics. The Mean Squared Error (MSE) measures the average squared difference between predicted and true copy number values, with WES data serving as the baseline. Cosine Similarity evaluates the similarity between CNV profiles by comparing them as high-dimensional vectors, with values closer to 1 indicating higher similarity. Euclidean Distance quantifies the straight-line distance between predicted and true CNV values, while Manhattan Distance calculates the absolute distance between these values. The Silhouette Score assesses clustering performance by measuring intra-cluster cohesion and inter-cluster separation, with higher values (closer to 1) indicating well-defined and well-separated clusters (Supplementary Note 4).

### Spatial domain identification

Copy number features were clustered using the Leiden algorithm[32], which optimizing modularity to identify communities in graphs. The modularity function is defined as:

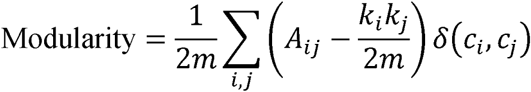

where *A_ij_* is the adjacency matrix, *k_i_* and *k_j_* are the degrees of nodes *i* and *j*, *m* is the total number of edges, and *δ*(*c_i_*, *c_j_*) equals 1 if nodes i and j are in the same community, and 0 otherwise. This approach segments the tissue into distinct spatial domains, corresponding to different tumor clones.

### Tumor Evolution Pattern Inference

The ultimate goal of SCOIGET is to infer tumor evolution patterns. High-dimensional embeddings capture both spatial and genomic features of the tumor microenvironment, providing a comprehensive representation of tumor dynamics. Tumor clones are identified based on clustering results, and their evolutionary relationships are analyzed through Partition-based Graph Abstraction (PAGA)[33], which constructs a phylogenetic tree to visualize the progression and diversification of tumor clones. This evolutionary tree enhances our understanding of tumor development and metastasis.

### Survival analysis

To evaluate the prognostic significance of differentially expressed genes identified among subclones by the SCOIGET algorithm, we performed a survival analysis using the GEPIA2 web server[34] (http://gepia2.cancer-pku.cn/) with data from the TCGA-COAD (Colon Adenocarcinoma) cohort. Specifically, the genes deemed differentially expressed between subclones were used to stratify patients into high- and low-expression groups, based on the median expression value (Group Cutoff = Median). Overall Survival (OS) was chosen as the clinical endpoint. GEPIA2 then generated Kaplan–Meier survival curves for each gene, and statistical significance was assessed using the log-rank test.

## Results

### Overview of SCOIGET

We introduce SCOIGET, a novel framework designed to infer copy number variations (CNVs) from spatial omics data by leveraging graph neural networks (GNNs) with graph attention layers (GAT). By integrating spatial gene expression profiles with multi-omics features, SCOIGET constructs a comprehensive spatial map of tumor heterogeneity, offering valuable insights into the underlying evolutionary processes.

The architecture of SCOIGET consists of several key components. The input layer combines transcriptomic profiles with spatial coordinates to build a spatial graph, encoding the neighborhood relationships between cells or tissue regions **(Fig. 1A)**. This is followed by the core model, which employs an encoder-decoder structure. The encoder uses GAT layers to extract meaningful features from the graph, while the decoder reconstructs these features, preserving both spatial and transcriptional information. In addition, the CNEncoder integrates a Hidden Markov Model (HMM) to detect CNV segments, capturing chromosomal alterations that are critical for understanding tumor evolution **(Fig. 1B)**.

**Figure 1.**
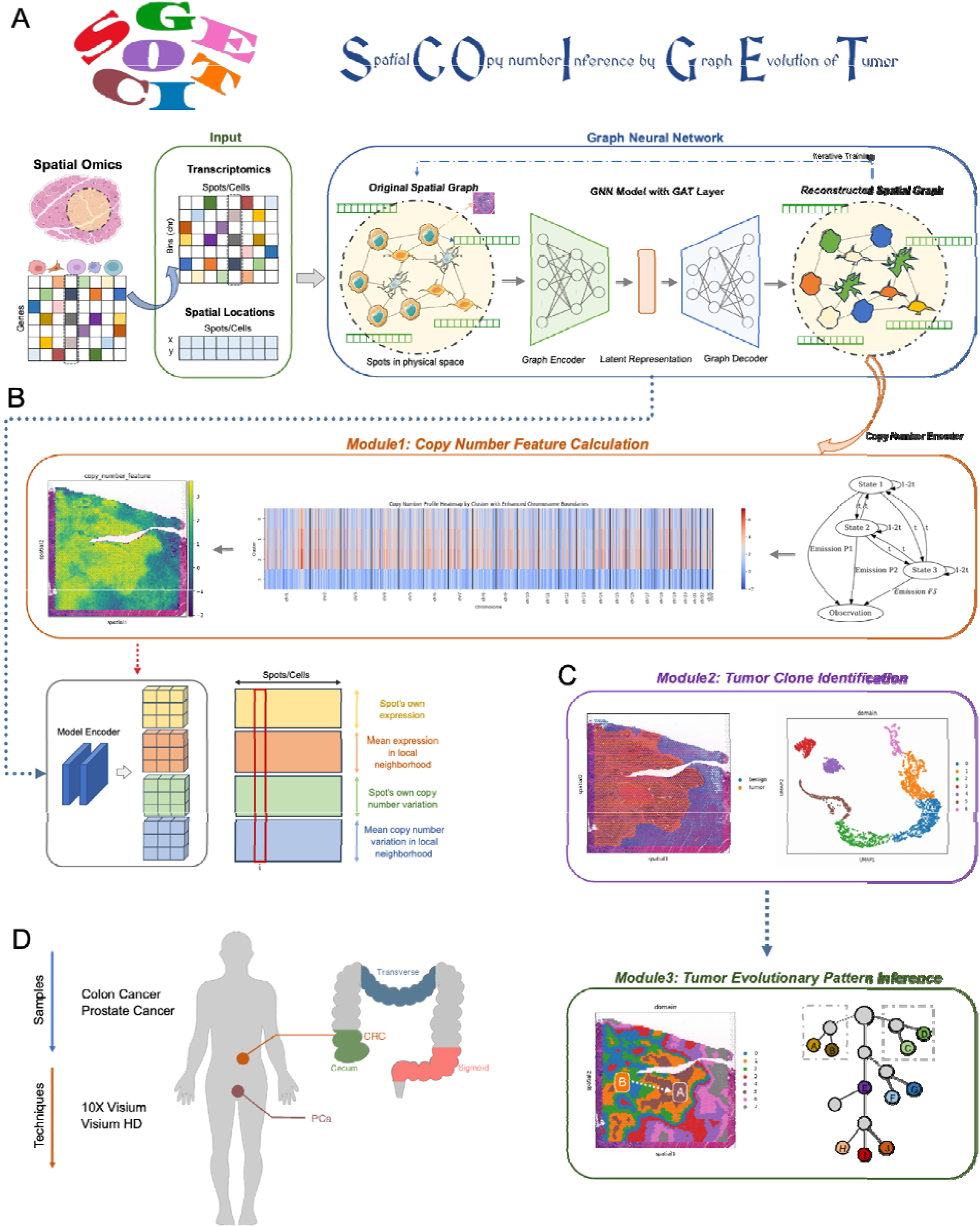
Overview of the SCOIGET framework. (A) **Model architecture.** The SCOIGET framework is built on a graph neural network (GNN) with Graph Attention Network (GAT) layers. It integrates spatial transcriptomics data with genomic features to infer copy number variations (CNVs) and detect tumor heterogeneity. The architecture includes an encoder that learns spatial-transcriptomic-genomic representations, a decoder that reconstructs input features, and a CNEncoder that estimates CNVs using a Hidden Markov Model (HMM). (B) **Module 1: Copy number feature calculation.** This module computes CNV features by predicting copy number profiles from spatial transcriptomics data, using the CNEncoder and HMM to identify and refine CNVs across the tissue. Spatial smoothing is applied to improve the localization of CNV segments. (C) **Module 2: Tumor clone identification and Module 3: Tumor evolutionary pattern inference.** Module 2 identifies spatial clones by clustering regions with similar genomic and transcriptomic characteristics. Module 3 infers tumor evolutionary patterns, reconstructing clonal dynamics and spatial-temporal tumor progression based on CNV profiles and clustering results. (D) **Dataset information.** The datasets used for evaluation include four spatial transcriptomics cohorts: three colorectal cancer (CRC) datasets (HTAN-WUSTL, Cell 2023, and 10x Genomics) and one prostate cancer (PCa) dataset (Nature 2022). These datasets were used for model validation and case studies, with matched WES data providing the gold standard for CNV estimation. Detailed dataset information is provided in Supplementary Table 1.

SCOIGET supports a variety of downstream analyses through three main modules. The first module calculates CNV features, providing quantitative measures of genomic alterations (Fig. 1B). The second module identifies spatial clones by grouping tissue subpopulations with similar genomic and transcriptomic profiles. Finally, the third module infers tumor evolutionary patterns, enabling the reconstruction of clonal dynamics and providing insights into the spatial and temporal progression of the tumor **(Fig. 1C)**. Together, these modules position SCOIGET as a powerful and versatile tool for exploring spatial tumor heterogeneity and advancing personalized oncology strategies.

### SCOIGET integrates spatial omics features within a unified framework and accurately detects copy number features spatially

To assess the accuracy of CNV inference, we evaluated SCOIGET on eight colorectal cancer (CRC) samples from the HTAN-WUSTL dataset (including HT260C1, HT112C1 [U1, U2], and HT225C1 [U1–U5]). The CNV profiles inferred by SCOIGET were benchmarked against matched whole-exome sequencing (WES) data and compared with four established CNV inference methods (InferCNV, CopyVAE, CopyKAT, and SCEVAN).

As shown in **Fig. 2A**, SCOIGET consistently outperformed alternative methods across several quantitative metrics, including mean squared error (MSE), cosine similarity, Euclidean distance, and Manhattan distance. For example, in sample HT260C1, SCOIGET achieved an MSE of 1.0764, a cosine similarity of 0.903, and both average Manhattan and Euclidean distances of 0.949. In contrast, InferCNV yielded an MSE of 1.6386, a cosine similarity of 0.8525, and distance metrics exceeding 1.15. Similarly, in samples HT112C1-U1 and HT112C1-U2, SCOIGET reported MSE values of 0.1098 and 0.1149, with cosine similarities of 0.9536 and 0.9598, respectively—significantly better than those of the competing methods. Overall, these statistical evaluations indicate that SCOIGET reduces error metrics by 30–80% while achieving superior pattern congruence.

**Figure 2.**
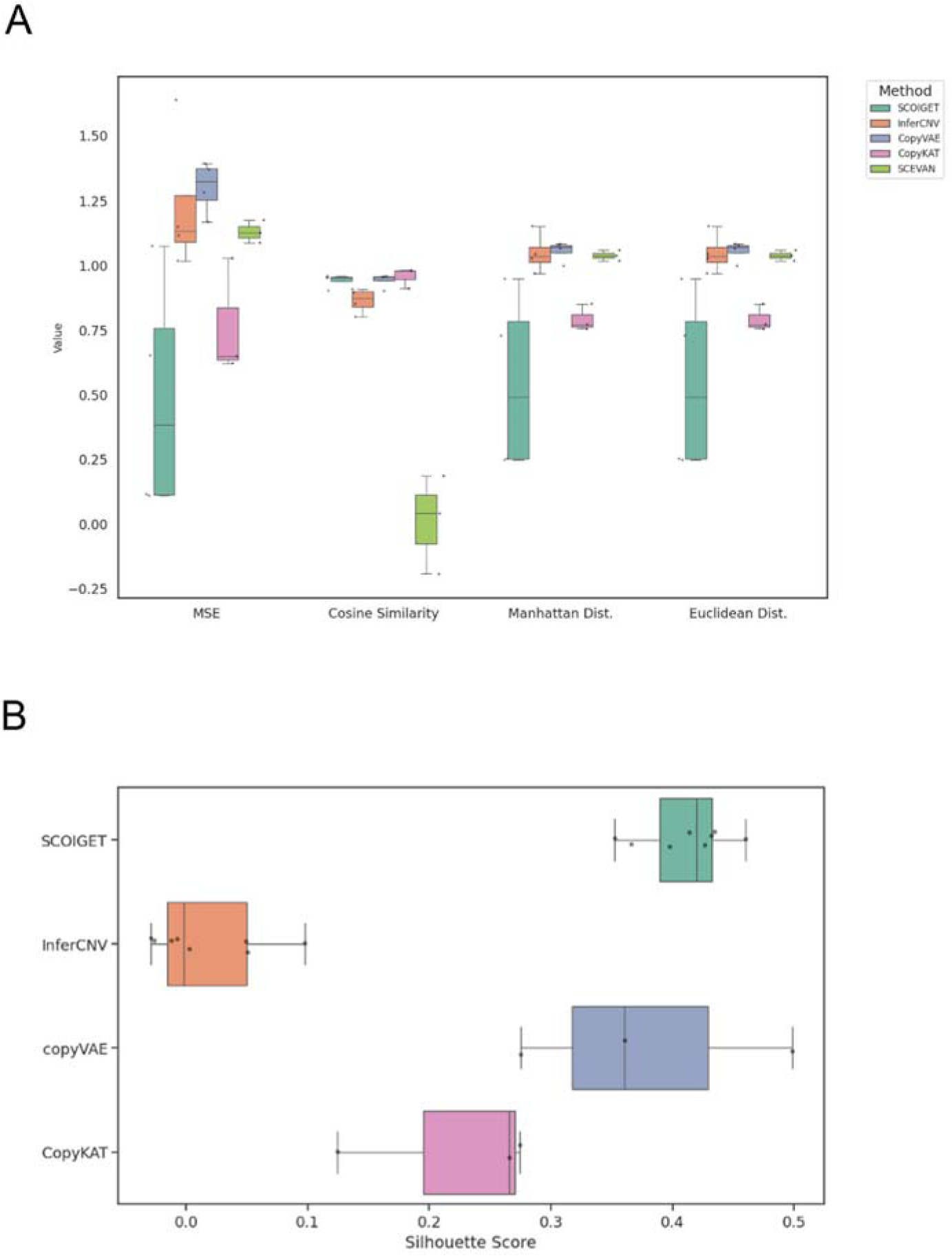
Model accuracy and comparison. (A) **Performance metrics for CNV inference.** The accuracy of SCOIGET’s CNV predictions was assessed using four metrics: Mean Squared Error (MSE), Cosine Similarity, Euclidean Distance, and Manhattan Distance. MSE quantifies the average squared difference between predicted and true CNV values (using WES data as the baseline). Cosine Similarity measures the similarity between predicted and true CNV profiles as high-dimensional vectors, with values closer to 1 indicating higher similarity. Euclidean and Manhattan distances capture the straight-line and absolute differences, respectively, between predicted and true values. SCOIGET outperformed other methods across all metrics, achieving the best overall performance. (B) **Clustering performance.** The Silhouette Score was used to evaluate clustering performance based on the CNV features inferred by SCOIGET. Higher Silhouette Scores indicate better-defined clusters with greater intra-cluster cohesion and inter-cluster separation. SCOIGET-derived CNV features resulted in the highest Silhouette Score, demonstrating superior clustering performance compared to other methods.

In addition to its high inference accuracy, the spatial CNV features derived from SCOIGET were also evaluated for downstream clustering applications. Using the Silhouette Score as a measure of clustering quality, SCOIGET-derived features consistently produced the highest scores— ranging from 0.3525 to 0.4599 in HT225C1 sub-samples, 0.4316 in HT260C1, 0.3973 in HT112C1-U1, and 0.4137 in HT112C1-U2. In comparison, clusters derived from InferCNV exhibited scores ranging from –0.0293 to 0.0976, while CopyVAE showed variable performance across samples (**Fig. 2B**). These results demonstrate that SCOIGET not only enhances the accuracy of CNV estimation but also yields more coherent and well-defined clusters.

Collectively, these findings highlight the effectiveness of SCOIGET in integrating spatial information for CNV inference, leading to enhanced concordance with independent genomic data and robust performance in downstream analyses. This integrated approach provides a powerful tool for elucidating tumor heterogeneity and advancing personalized oncology strategies.

### Inferring clonal evolution in colorectal cancer progression through spatial CNV analysis

To investigate tumor evolution, we applied SCOIGET to four 10x Visium CRC samples derived from the same patient’s cecum, including one G1-stage sample (6723_4, TVA subtype) and three G2-stage MSS samples (6723_1, 6723_2, 6723_3). Following integration of basic expression profiles with Harmony, SCOIGET generated spatial CNV scores that delineated tumor boundaries and highlighted malignant potential (**Fig. 3A**). Leiden clustering of these CNV features identified spatially distinct domains corresponding to the tumor’s clonal architecture (**Fig. 3C**). High-CNV-score domains (1, 2, 3, 8) were centrally localized, while intermediate-CNV domains (0, 5, 7, 9, 10, 12, 13) and low-CNV-score domains (4, 6, 11) occupied the periphery. These domains were associated with four subclones: cloneA (domain2), cloneB (domain8), cloneC (domain1), and cloneD (domain3). CloneA and cloneB were exclusive to the cancerous stage, cloneC spanned both precancerous and cancerous phases, and cloneD marked a transitional boundary (**Fig. 3H**).

**Figure 3.**
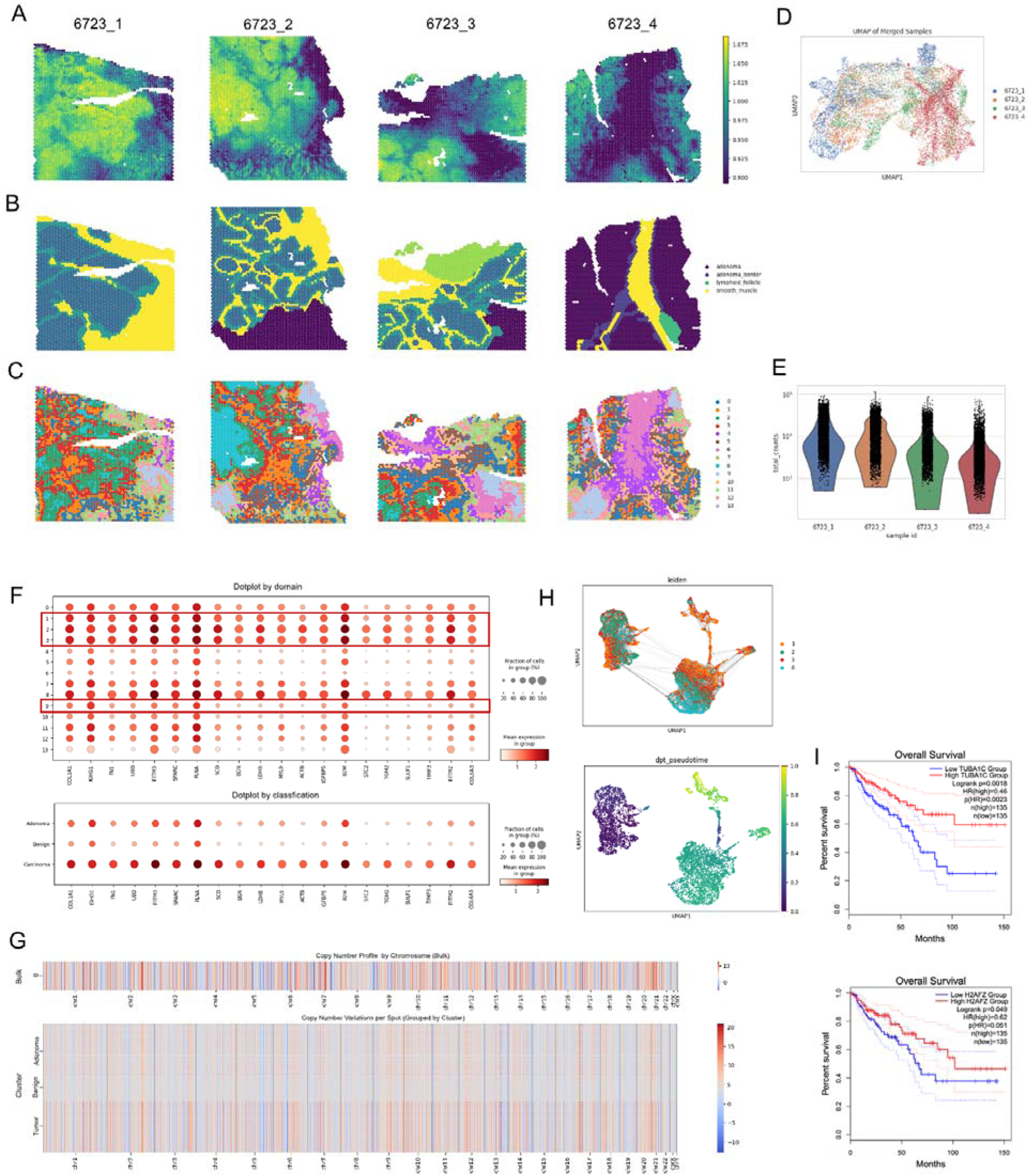
Clonal Evolution in Colorectal Cancer Inferred through Spatial CNV Analysis. (A) **Spatial CNVs Inferred by SCOIGET.** Visualization of copy number variations across the tumor, highlighting regions with distinct CNV patterns. (B) **Pathologist Annotations.** Histological annotations based on H&E-stained images used to validate CNV-based subclones. (C) **Leiden Clustering of CNV Features.** Clustering results showing spatially distinct domains with varying CNV scores. (D) **UMAP Visualization of Patient Samples.** Four patient samples (6723_1, 6723_2, 6723_3 from G2 stage [cancer], and 6723_4 from G1 stage [precancer]) visualized in UMAP space after grouping by tumor stage. (E) **Total Gene Expression Counts.** Bar plot illustrating the total counts of gene expression across the four samples. (F) **Differential Gene Expression Dotplot.** Dotplot showing differentially expressed genes in the 14 domains identified in panel (C), grouped into benign, adenoma, and carcinoma categories. (G) **Chromosome-Level CNV Heatmaps.** Heatmaps displaying copy number variations segmented by chromosomes for bulk tumor samples and subgrouped domains (benign, adenoma, carcinoma). (H) **Identification of Clones and Clonal Trajectories.** Four distinct clones (cloneA, cloneB, cloneC, cloneD) identified within carcinoma-associated domains, with UMAP projections and clonal trajectories inferred using PAGA analysis. CloneA and cloneB appear only in the cancerous stage, cloneC spans both G1 and G2 stages, and cloneD marks the transitional zone. (I) **Survival Analysis of Differential Genes.** Survival curves for differential genes TUBA1C and H2AFZ in cloneB, showing significant prognostic differences in CRC patient survival based on TCGA data.

Pathologist annotations of HE-stained images validated the CNV-based classifications (**Fig. 3B**). UMAP visualization of the four samples by tumor stage confirmed the spatial segregation of cancerous and precancerous regions (**Fig. 3D**), while total gene expression counts across these stages highlighted distinct expression patterns (**Fig. 3E**). Differentially expressed genes among the 14 domains reinforced the CNV-based classifications, with cancerous regions showing elevated expression of CRC-associated markers (**Fig. 3F**). Chromosome-level CNV heatmaps further delineated alterations across tumor subtypes, distinguishing benign, adenoma, and carcinoma regions (**Fig. 3G**).

By integrating chromosome-level CNV profiles and assessing clonal trajectories, SCOIGET pinpointed cloneC as pivotal in transitioning from G1 to G2 stages, suggesting its involvement in early tumor progression (**Fig. 3H**). Furthermore, survival analysis of cloneB-specific genes, such as TUBA1C and H2AFZ, revealed significant prognostic differences in CRC patients based on TCGA data (**Fig. 3I**). TUBA1C, which encodes a tubulin alpha chain essential for microtubule dynamics, is overexpressed in colorectal and other cancers. Its elevated expression is associated with poor prognosis, tumor progression, and modulation of the tumor microenvironment, suggesting its potential as both a prognostic biomarker and therapeutic target[35,36]. Meanwhile, H2AFZ, a histone variant involved in chromatin remodeling and transcriptional regulation, plyas a critical role in tumorgenesis. Its overexpression has been stronly associated with enhanced tumor aggressiveness epithelial–mesenchymal transition (EMT), and poor survival outcomes across various cancers, underscoring its importance as a prognostic indicator[37,38].

These findings reinforce the biological significance of these markers and highlight how SCOIGET’s robust CNV estimates can direct downstream functional analyses, ultimately enhancing our understanding of tumor progression and guiding targeted, precision oncology strategies.

### SCOIGET reveals tumor evolution patterns in prostate cancer

To evaluate model’s generalizability, we applied SCOIGET to four Visium PCa samples collected from distinct regions of the same patient. SCOIGET effectively inferred spatial CNV features, identifying both shared and region-specific genomic alterations (**Fig. 4A**). Leiden clustering revealed spatial domains corresponding to Gleason Grades (GG1, GG2, GG4), border regions, and benign areas, aligning with histological classifications and revealing finer structures(**Fig. 4C**). These clustering results were validated by pathologist annotations from HE-stained images, demonstrating strong concordance (**Fig. 4B**).

**Figure 4.**
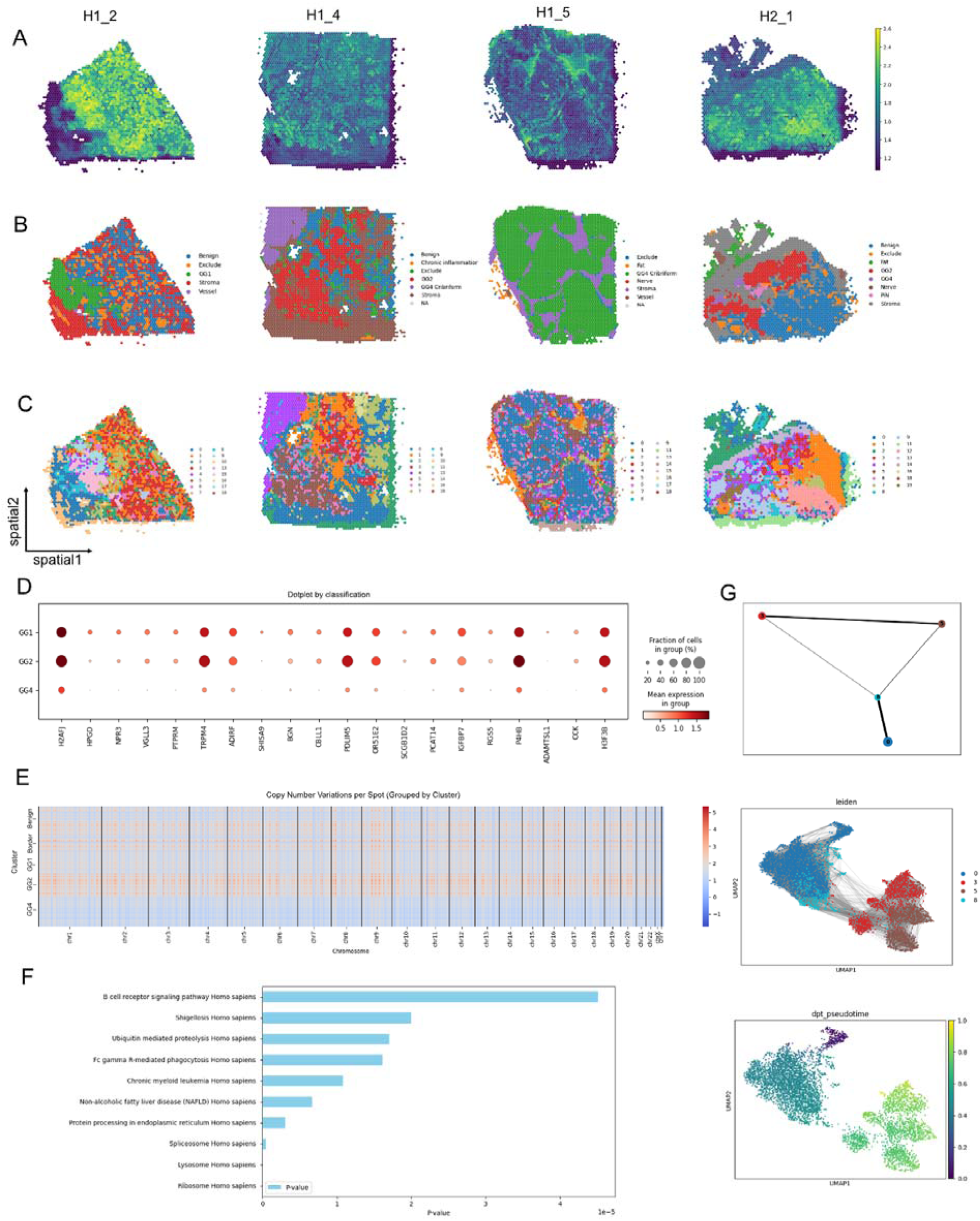
Tumor Evolution Patterns in Prostate Cancer Revealed by SCOIGET. (A) **Spatial CNVs inferred by SCOIGET.** Visualization of copy number variations across tumor regions, capturing distinct CNV patterns in prostate cancer samples. (B) **Pathologist annotations.** HE-stained histological images annotated by pathologists, validating SCOIGET-derived spatial CNV clusters. (C) **Leiden clustering of CNV features.** Spatial domains identified based on CNV features, corresponding to Gleason Grades (GG1, GG2, GG4), border, and benign regions. (D) **Differential gene expression dotplot.** Dotplot showing distinct gene expression profiles across tumor grades, highlighting key genes associated with tumor progression. (E) **Chromosome-level CNV heatmaps.** Heatmaps of copy number variations grouped by chromosome, demonstrating genomic heterogeneity among different Gleason Grades and benign regions. (F) **Pathway enrichment analysis.** Bar plot of enriched pathways among differentially expressed genes across tumor clusters, emphasizing pathways linked to immune regulation and tumor progression. (G) **Clonal evolution trajectories inferred by PAGA.** PAGA-based analysis mapping tumor clonal evolution, identifying cluster 0 as the root node and illustrating the progression and divergence of tumor clones.

Further analyses illuminated molecular distinctions between tumor grades. Differentially expressed genes among clusters were enriched in pathways associated with tumor progression and immune modulation, including the B cell receptor signaling pathway, ubiquitin-mediated proteolysis, and chronic myeloid leukemia (**Fig. 4F**). Chromosome-level CNV heatmaps showcased unique and shared alterations across tumor grades, providing detailed insights into genomic heterogeneity (**Fig. 4E**).

Using PAGA, SCOIGET reconstructed clonal evolution trajectories, identifying cluster 0 as the root node and mapping the progression and divergence of tumor clones (**Fig. 4G**). These integrated analyses underscore SCOIGET’s utility in elucidating clonal dynamics and identifying biologically significant features, offering potential insights into therapeutic targets and prognostic biomarkers.

These findings affirm SCOIGET’s ability to resolve tumor evolution within heterogeneous tissue environments. By integrating spatial omics with CNV inference, SCOIGET provides a robust platform for investigating tumor heterogeneity, advancing biomarker discovery, and supporting precision oncology.

### Unveiling colorectal cancer heterogeneity at subcellular resolution

To further validate its versatility, we applied SCOIGET to subcellular-resolution spatial transcriptomics data from three CRC samples (p1, p2, and p5), each collected from a distinct patient. After annotating cell types using established marker genes, we distinguished malignant and benign regions to serve as a reference framework (**Fig. 5A, D, F**). SCOIGET-derived CNV features revealed a strong correspondence between spatially inferred genomic alterations and tumor microenvironment boundaries, effectively highlighting transitions between tumor and non-tumor areas (**Fig. 5B, E, G**).

**Figure 5.**
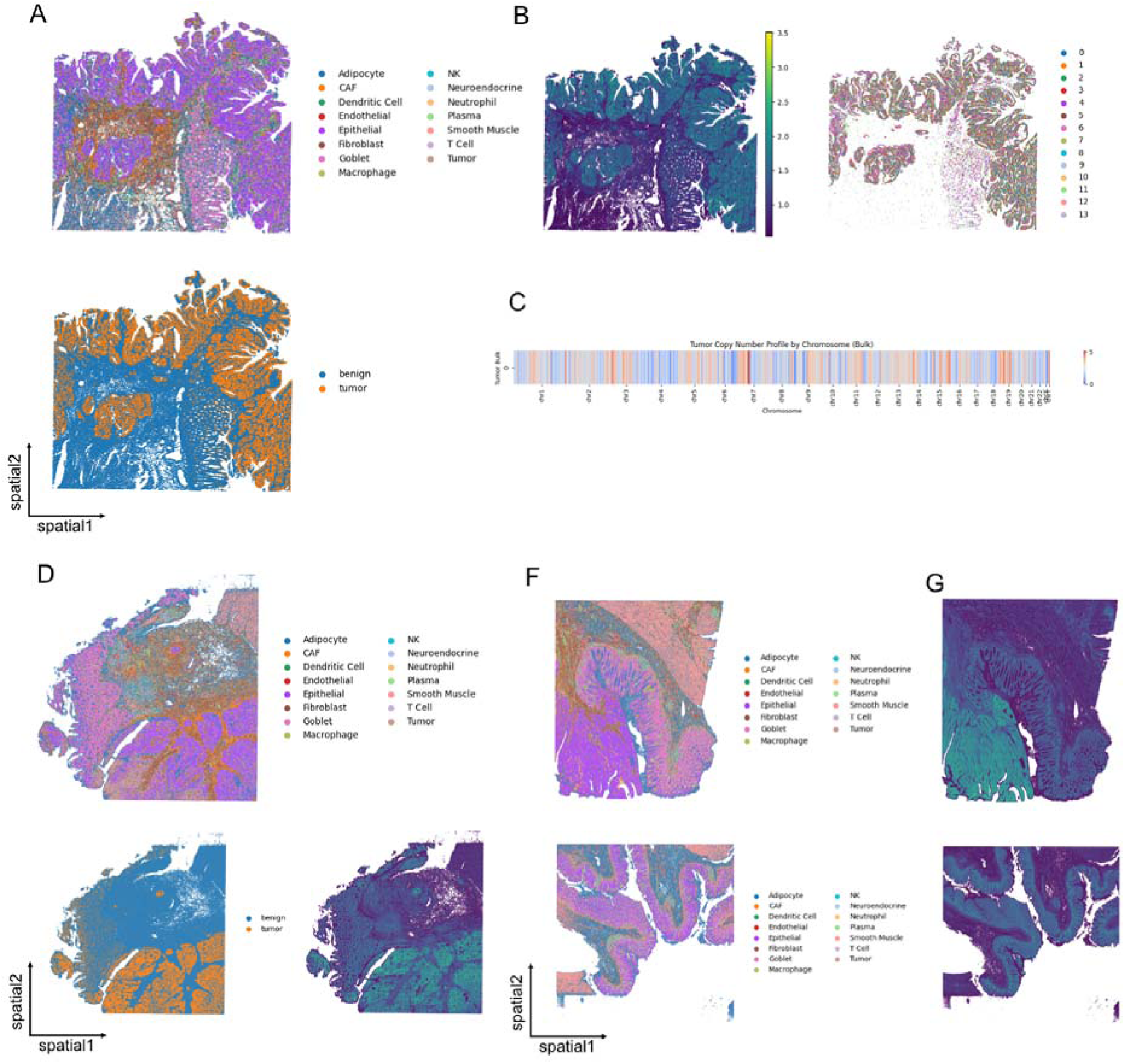
SCOIGET applied to subcellular-resolution CRC spatial omics data. **(A)** Annotation of cell types in the p2 sample, with malignant and benign regions distinguished based on gene expression profiles. **(B)** Spatial CNV inference results for p2, coupled with Leiden clustering of tumor regions. **(C)** Chromosome-level bulk tumor CNV heatmap showcasing inferred genomic alterations. **(D)** Cell type annotations in the p1 sample, highlighting malignant and benign areas. **(E)** Spatial CNV inference results for p1, revealing tumor boundaries and genomic transitions. **(F)** Cell type annotations in the p5 sample, which includes CRC and NAT (normal adjacent tissue) regions. **(G)** Spatial CNV inference results for p5, illustrating tumor and control region differences.

Chromosome-level CNV predictions at the bulk tumor level further supported these observations, showcasing consistent alterations across spatial regions (**Fig. 5C**). Notably, SCOIGET successfully identified subtle spatial patterns of genomic alterations, even in subcellular-resolution data, offering insights into the spatial and evolutionary heterogeneity of CRC.

These results highlight SCOIGET’s adaptability to high-resolution spatial omics platforms, enabling detailed exploration of tumor microenvironment dynamics and evolution at single-cell resolution. Its ability to capture fine-grained spatial evolutionary patterns underscores its potential for broader applications, offering new avenues for interpreting tumor heterogeneity and progression.

## Discussion

In this study, we proposed an innovative approach that integrates spatial information to infer spatial copy number variations (CNVs), applying the derived features to tasks like identifying tumor clones, analyzing clone differences, and inferring evolutionary trajectories in spatial transcriptomics. By incorporating spatial dimensions, SCOIGET provides a more nuanced understanding of tumor heterogeneity, effectively linking spatial and genomic information to reveal insights into tumor progression and microenvironmental interactions. Its versatility is demonstrated through successful application to colorectal cancer and prostate cancer, uncovering clonal architectures and evolutionary dynamics in distinct tumor types. Furthermore, we extended SCOIGET to subcellular-resolution spatial transcriptomics data, highlighting its adaptability to high-resolution platforms.

Despite its promising results, inferring CNVs solely from transcriptomic data may have inherent challenges. Although gene expression data has been widely used in cancer research, the absence of direct causal evidence between expression patterns and CNVs introduces potential inaccuracies. Moreover, relying on a single omics layer limits the exploration of additional molecular mechanisms, such as mutations, epigenetic alterations, and proteomic interactions, which are critical for a comprehensive understanding of tumor biology.

Future research should focus on integrating multi-omics data—such as genomics, epigenomics, and proteomics—to improve the robustness and accuracy of SCOIGET’s CNV predictions. Expanding spatial features to include tumor-immune and tumor-stromal interactions could further enhance its capability to delineate complex tumor microenvironments. SCOIGET’s demonstrated performance across different cancers and resolution levels underscores its potential as a transformative tool in cancer research, supporting biomarker discovery, therapeutic target identification, and personalized oncology strategies.

## Supplementary Data

Attached in the supplementary file.

## Data Availability

The study uses publicly available spatial transcriptomics data. CRC Visium data is retrieved from HTAN WUSTL and Vanderbilt Atlas by HTAN DCC Portal (https://data.humantumoratlas.org/). CRC VisiumHD data is retrieved from 10X genomics (https://www.10xgenomics.com/products/visium-hd-spatial-gene-expression/dataset-human-crc/). PCa Visium data is retrieved from Mendeley Data (https://doi.org/10.17632/svw96g68dv.1).

## Code Availability

The code is publicly available at https://github.com/YukiZH/SCOIGET under MIT license.

## Author’s Contributions

Y.J.Z. contributed to the experimental design, conceptualization, analysis, and biological interpretation. Y.T.Y. designed the model, while Y.K. assisted with conceptualization and biological interpretation. B.X.Z. contributed to testing and debugging the code. Y.J.Z. drafted the manuscript, and Y.T.Y., Y.K., B.X.Z., K.N., and H.L. critically revised it. K.N. and H.L. supervised the project. All authors reviewed and approved the final manuscript.

## Funding

This research is supported by the Science and Technology Commission of Shanghai Municipality (Grant No. 23JS1400700), Neil Shen’s SJTU Medical Research Fund, and the Science and Technology Innovation Key R&D Program of Chongqing (Grant No. CSTB2024TIAD-STX0006).

## Notes

### Competing Interest Statement

The authors have declared no competing interest.

